# Fast trimer statistics facilitate accurate decoding of large random DNA barcode sets even at large sequencing error rates

**DOI:** 10.1101/2022.07.02.498575

**Authors:** William H. Press

## Abstract

Predefined sets of short DNA sequences are commonly used as barcodes to identify individual biomolecules in pooled populations. Such use requires either sufficiently small DNA error rates, or else an error-correction methodology. Most existing DNA error-correcting codes (ECCs) correct only one or two errors per barcode in sets of typically ≲ 10^4^ barcodes. We here consider the use of *random* barcodes of sufficient length that they remain accurately decodable even with ≳ 6 errors and even at ~ 10% or 20% nucleotide error rates. We show that length ~ 34 nt is sufficient even with ≳ 10^6^ barcodes. The obvious objection to this scheme is that it requires comparing every read to every possible barcode by a slow Levenshtein or Needleman-Wunsch comparison. We show that several orders of magnitude speedup can be achieved by (i) a fast triage method that compares only trimer (three consecutive nucleotide) occurence statistics, precomputed in linear time for both reads and barcodes, and (ii) the massive parallelism available on today’s even commodity-grade GPUs. With 10^6^ barcodes of length 34 and 10% DNA errors (substitutions and indels) we achieve in simulation 99.9% precision (decode accuracy) with 98.8% recall (read acceptance rate). Similarly high precision with somewhat smaller recall is achievable even with 20% DNA errors. The amortized computation cost on a commodity workstation with two GPUs (2022 capability and price) is estimated as between US$ 0.15 and US$ 0.60 per million decoded reads.

## 1 Introduction

The use of DNA barcode libraries to identify tagged individual biomolecules in pooled populations has become an essential tool for today’s massively parallel biomedical experiments. Barcodes find use in gene synthesis[1, 2], antibody screens [3, 4], drug discovery via tagged chemical libraries [5–7], and many other applications [8–14], including their potential use in schemes for engineered DNA data storage [15, 16]. For some applications, barcodes must function robustly in experimental situations subject to significant error rates (that is, the unintended occurrence of nucleotide substitutions, insertions, and deletions). Errors may be introduced during barcode synthesis, the processes of the experiment, the final sequencing, or all of these [15]. Errors in barcode synthesis (“wrong barcodes”) are particularly troublesome, because they create errors that persist at any depth of final sequencing.

“Next-Generation Sequencing” (NGS), as exemplified in Illumna technology [17], has relatively short read lengths (200-300 nt), but also relatively small error rates (10^−3^–10^−4^ per nt). In this regime, barcodes need to be short (≲ 20 nt), but they need only modest (if any) error-correction capability. In other words, barcodes as ultimately sequenced can be assumed to have at most one or two errors, allowing the use of repurposed mathematical error-correcting codes (ECCs) [18, 19], sometimes [20–24], but not always [25, 26], with the necessary extensions to account for insertion and deletion errors (“indels”). By a “mathematical” ECC, we mean a set of codewords and also an algorithm for recovering an original codeword from its garbled version, specifically without needing to compare every potentially garbled read to every known possible codeword in the library. (We use the terms “barcode” and “codeword” almost interchangeably, the former being the physical manifestation in DNA of the mathematical latter.)

Useful mathematical ECCs for use with NGS have in practice been limited to libraries with no more than tens of thousands of unique codewords. While any code should ideally signal a reject (“erasure”) rather than return a wrong identification if the garbled word has more errors than the ECC can handle, this in general cannot be mathematically assured if the number of errors is not strictly bounded [19].

Hawkins et al.’s “FREE” barcodes [27] overcame some of these limitations with a direct approach: Libraries of pairwise dissimilar codewords were constructed by comparing each proposed new codeword to all previously accepted ones, a slow, but one-time, process. A novel similarity measure was designed to be tolerant of indels that could produce garbled barcodes with unknown, altered lengths. Advantageously, codewords could be constrained to have balanced GC content, minimal homopolymer runs, reduced hairpin propensity, or any other experimentally motivated constraints. In the FREE scheme, garbled codewords are decoded by table lookup into a very large table containing not only the codewords, but also all of their possible single- or double-error garbles. This is very fast, but requires very large computer memory. Practically, this scheme achieves single-error–correcting codes of 16-nt length, with 1.6 × 10^6^ barcodes, or double-error–correcting codes of 17-nt length with 23,000 codes.

In recent years, third-generation sequencing (also known as TGS or long-read sequencing), in variants developed by Pacific Biosciences and Oxford Nanopore [28] has changed the landscape. TGS is capable of very long reads, *>* 10^4^ nt, so barcode length is of small consequence. However, read error rates may be as high as ∼ 10% [29] Such error rates render virtually useless single- and double-error correcting barcode libraries of useful size. In the most favorable case of independent random errors, three or more errors can occur frequently; burst errors such as stuttering or repeated deletions only make things worse.

This paper explores a possible solution via the use of *random* barcodes (“randomers”) [30] that is, barcode libraries of any desired size (≳ 10^6^, for example) whose codewords are approximately uniformly random, as generated by computer, with constraints of GC content, homopolymers, etc., easily imposed. Like “designed” barcodes, random barcodes would be synthesized in defined oligo pools, but with the difference that the number of pools could be as large as desired. There are two obvious, immediate objections to this scheme that must be overcome: 1. How can we avoid too-similar pairs of codewords in the library, so that the garbles of one are not mistakenly decoded as the other? 2. How can we avoid the impractical all-to-all in-silico comparison of every read to every codeword in the library?

The answers are unexpectedly simple: 1. We use barcodes of length sufficient to make near-collisions statistically unlikely to any desired degree. To implement this, we below investigate the statistics of such near-collisions. 2. Instead of rejecting all-against-all brute force comparison, we embrace it. Below, we will describe a novel, fast computational technique that characterizes codewords by their overlapping trimers (three-nucleotide sequences), both trimer presence versus absence and the order of those present. We show in particular that these techniques can run with massive parallelism on commodity graphics processing units (GPUs) and that cloud GPU availability [31],[32] makes such all-against-all comparisons practical at low cost and with reasonable throughput.

## 2 Materials and Methods

### 2.1 Distance Measures

Given a set of barcode codewords, and given a garbled barcode read (possibly, because of indels, prefixed or suffixed by spurious nucleotides), by definition the best decode we can do is to assign the read to its most probable codeword—or to declare it an erasure that cannot be reliably so assigned. “Most probable” implies an accurate statistical characterization of all the processes that produce errors, in practice rarely available [33]. So, any practical procedure involves choosing a surrogate, a distance measure between the two strings that at least approximates (a monotonic function of) *P* (*R*|*C*), the probability of a garbled read *R* given the true codeword *C*.

A gold standard for such an approximation, is the Needleman-Wunsch [34] alignment distance between the strings, with the skew, substitution, insertion, and deletion penalties set to the negative log-probabilities of their respective occurrence in an experimentally validated error model. To the degree that errors are independent, the distance so obtained is the negative log-probability of the most probable single path from codeword to read. Note that even this gold standard is not exact, because (i) the implied model of independent and identically distributed (i.i.d.) errors is surely not right in detail, and (ii) the probability *P* (*R*|*C*) is actually a sum over all possible paths, not the single most probable path.

Levenshtein distance (also called edit distance) [22] is a kind of silver standard, not as good as Needleman-Wunsch, but also not dependent on knowing error probabilities. Levenshtein distance is identical to Needleman-Wunsch when the skew, substitution, insertion, and deletion penalties all set to the same constant value (without loss of generality the value 1). In the remainder of this paper, we will use Levenshtein distance exclusively. However, all of the algorithms developed (and all of the implementing computer code) is designed to allow arbitrary penalties, hence the easy generalization to Needleman-Wunsch.

### 2.2 Levenshtein Distance Distribution of Random Strings

If there were no indels, then the Levenshtein distance between two random strings of the same length would be their Hamming distance, with an easily calculated binomial probability distribution (for independent errors). With indels, the distribution of Levenshtein distances between two random strings is a famously unsolved problem, closely related to the better-known unsolved problem of longest-common subsequences [35]. While it is known that for asymptotically long strings the mean distance scales as a constant *γ*_*c*_ times string length (hardly a surprise, given that the errors are local), *γ*_*c*_, termed the Chvátal-Sankoff constant [36] is not known, though it is conjectured to be 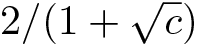, where *c* is the alphabet size (for us, 4). Beyond this mean, virtually nothing is known about the distribution of distances, although there is a conjectured connection to so-called Tracy-Widom distributions [37].

While little is known analytically, simulation is straightforward. Supporting Information S1 (text) describes how one-to-many Levenshtein distances can be parallelized on a GPU, allowing the calculation of *>* 10^8^ distances on a single-headed desktop machine in minutes. Figure 1 shows the results of such a simulation.

**Figure 1.**
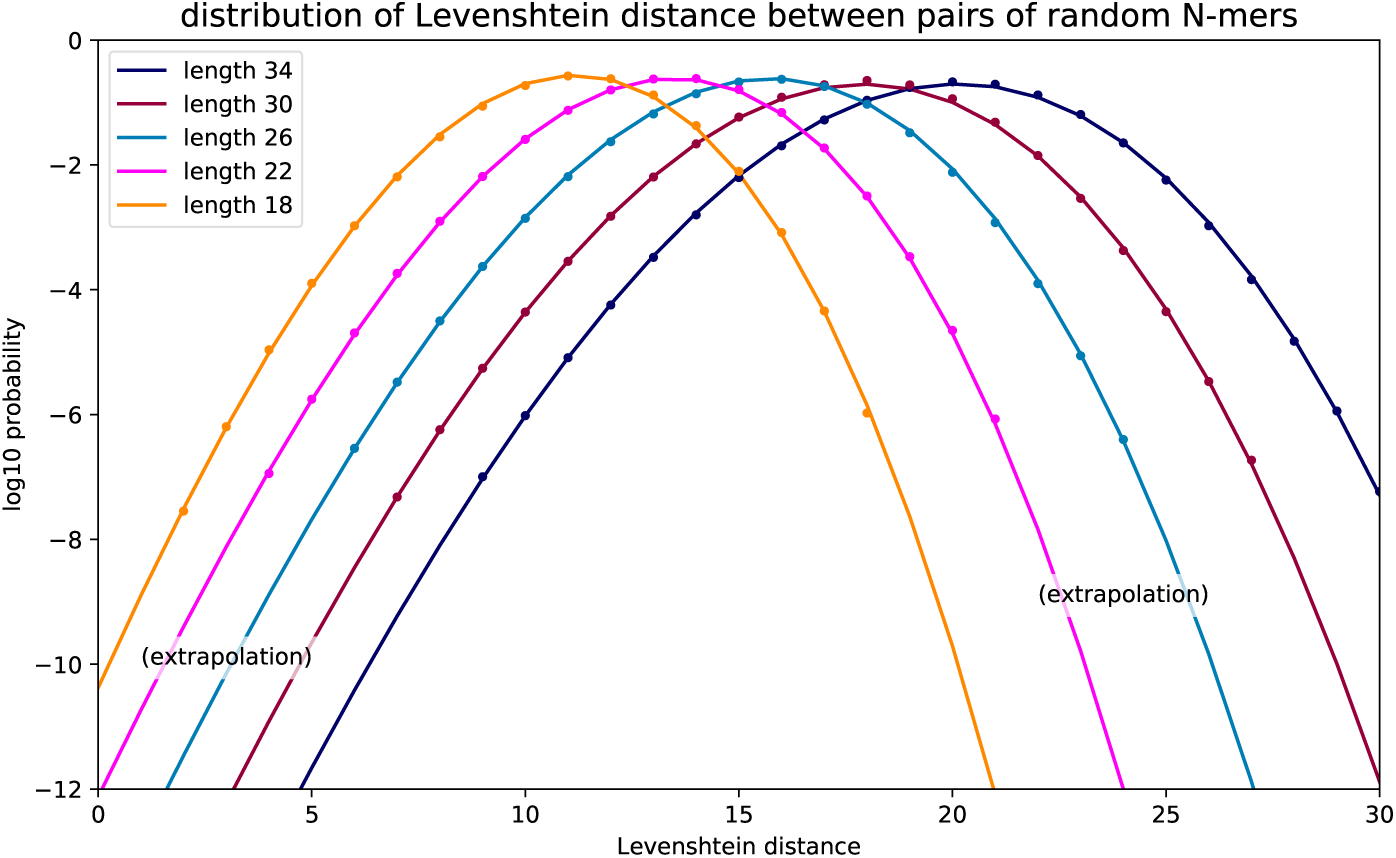
Probability distribution of Levenshtein distances. Random oligomers of lengths 18, 22, 26, 30, and 34 are generated and random pairwise Levenshtein distances are calculated. Dots show the results. The curves are a bivariate polynomial fit (in log-space) to all the dots simultaneously. The distributions are non-Gaussian in their tail, the curves deviating from parabolas slightly but significantly.

We are concerned about the extreme left-hand tails of the distributions, where a garbled read from one codeword might end up by unlucky chance close to another in a large set of barcodes. In this regime, direct sampling is impractical, but we can use the polynomial extrapolations (in log-space) shown in Figure 1. Their uncertainty at probability 10^*−*12^ is likely ≲ 1 in Levenshtein distance, as estimated by robustness of the curves as details of the fitting procedure are varied. Supporting Information S2 gives details of the polynomial fit shown in the figure.

### 2.3 Distribution of Closest Non-Causal Distance to a Set of N Codewords

Given a set of *N* random codewords {*C*} of length *M* from which a given read *R* does *not* derive, what is the probability *P* (*L*) that *R*’s smallest Levenshtein distance to the set is *L*? We may assume that *R* is itself (close to) random, because it derives from errors on a random starting point, its true codeword. Given one of the distributions in Figure 1, which we now denote *p*(*L*|*M*), this is a straightforward calculation in extreme value theory [38]: The probability *P* (0) is the cumulative Poisson probability of one or more zero distances when the mean number is *Np*(0|*M*),

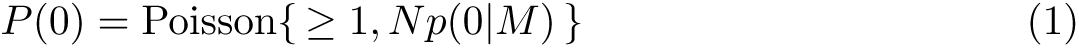

Then, recursively,

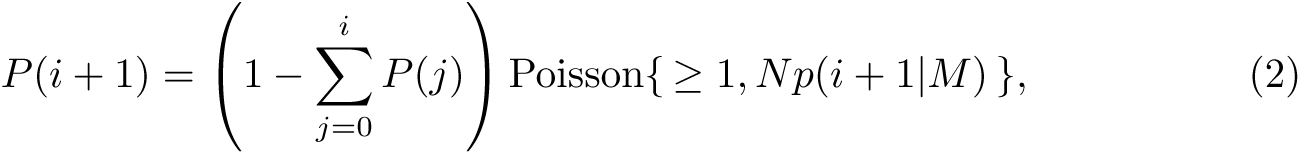

where the term in parentheses is the remaining probability to be allocated, and the Poisson cumulative distribution function is the probability of allocating it to the value *i* + 1. Equation 2 is easily computed numerically and is shown for the case of *M* = 34 nt in Figure 2 The values near the peaks seem haphazard due to discreteness effects, but are accurately shown.

The figure also plots the cumulative distribution functions for binomial deviates with parameters 34 (the codeword length) and probabilities 0.05, 0.10, and 0.20. These model, at least crudely, the Levenshtein distances to be expected in the causal case of comparison to the *correct* codeword. That fact that some orders of magnitude of vertical white space lie between each green curve and at least one other-colored curve points the way forward: By picking an appropriate threshold Levenshtein distance *T*, calling as decodes all reads with ≤ *T* and as erasures all reads with *> T*, we may hope to achieve both very high accuracy (high precision) on decodes and a very low erasure rate (high recall). The figure demonstrates this in an approximate, but relatively model-independent way. In Results, we will explore a more accurate, detailed model and, importantly, will give a procedure for choosing *T* based on observed data.

Supporting Information S3 shows figures analogous to Figure 2 for the cases of other codeword lengths, *M* = 30, 26, 22, and 18 nt.

**Figure 2.**
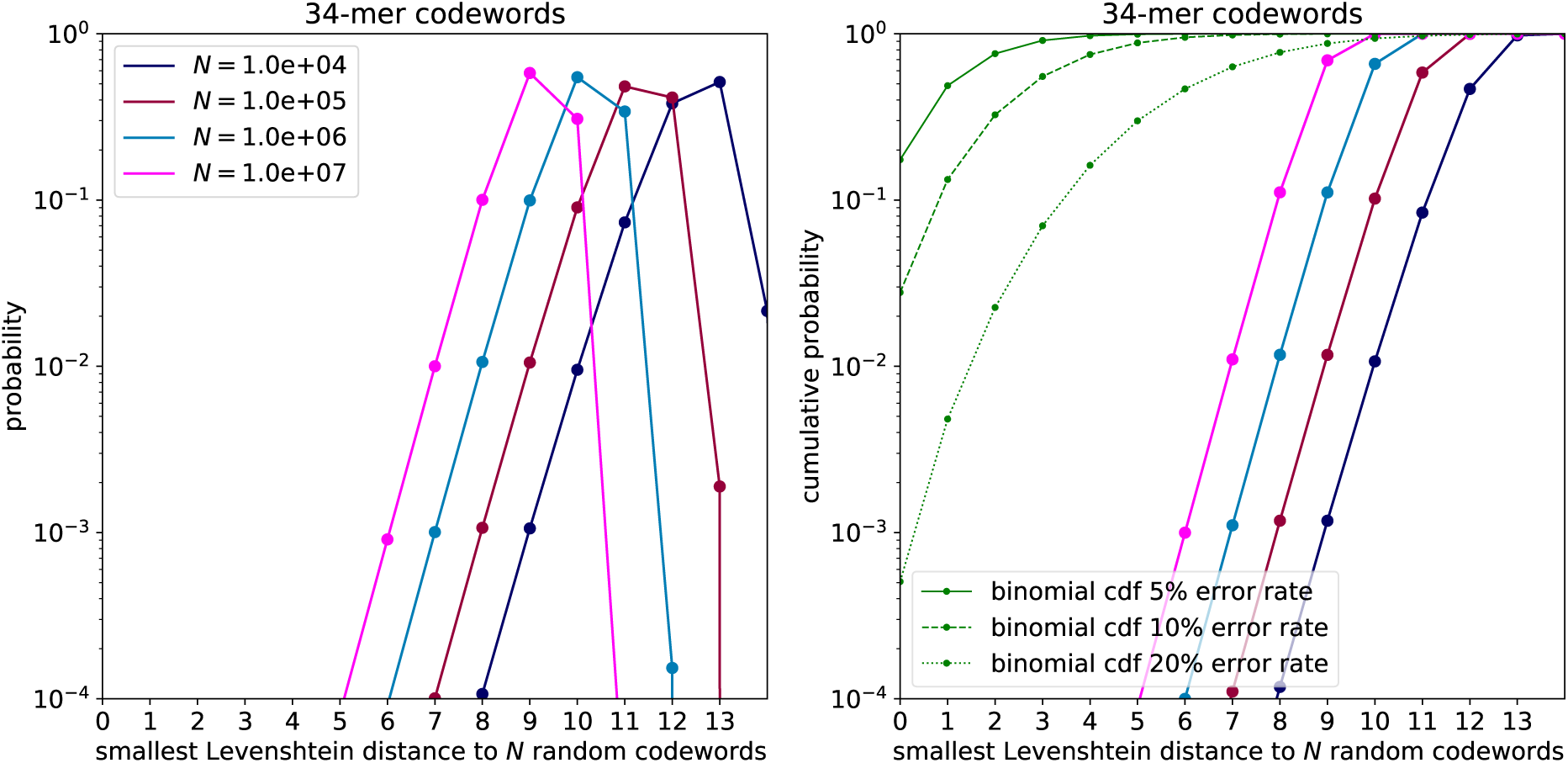
Probability of smallest distance to a set of *N* 34 nucleotide random codewords. Left: Probability mass function. The larger is *N*, the smaller is the expected distance to a given garbled read by chance. Right: Cumulative distribution function. Also shown as thin green lines are cumulative binomial probabilities for the number of errors in a garbled 34-mer for the large error rates (per nucleotide) 5%, 10% and 20%.

### 2.4 Three-parameter Poisson Error Model for Substitutions, Insertions, and Deletions

An error model for a *M* -mer barcode set can be described by three parameters *p*_*sub*_, *p*_*ins*_, and *p*_*del*_, respectively the probabilities per nucleotide of a substitution, insertion, or deletion error. Formally, we need to be more precise: the different types of errors can interact; and, insertion and deletion errors change the length of the string. Among various equally good possibilities, for purposes of this paper we adopt the following error-generation model, the steps to be executed in the order listed.

- Start with a codeword string in {*a, c, g, t*} of length *M*, indexed as 0 … *M −* 1.
- *Substitutions*. Draw a deviate 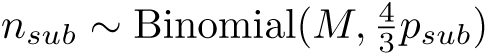. The factor 4*/*3 corrects for the fact that 1*/*4 of substitutions will substitute an unchanged nucleotide. Draw (with replacement) *n*_*sub*_ indices in the uniform distribution *U* (0, …, *M* − 1). Draw (with replacement) *n*_*sub*_ values uniformly in the nucleotides {*a, c, g, t*}. Substitute the values at the indices. (Note that indices may collide, in which case only one of the corresponding values will “win” the substitution, it doesn’t matter which.)
- *Deletions*. Draw a deviate *n*_*del*_ *∼* Binomial(*M, p*_*del*_). Draw (with replacement) *n*_*del*_ indices in the uniform distribution *U* (0, …, *M* − 1). Delete the nucleotides at those positions. Here, colliding indices delete the same position only once. Call the new string length *M* ′ *≥ M − n*_*del*_ with equality in the case of no collisions.
- *Insertions*. Draw a deviate *n*_*ins*_ ∼ Binomial(*M* ′, *p*_*ins*_). Draw (with replacement) *n*_*ins*_ indices in the uniform distribution *U* (0, …, *M* ′). Draw (with replacement) *n*_*ins*_ values uniformly in the nucleotides {*a, c, g, t*}. Insert each value before the original index position (or, for index *M* ′, after the last character). Here, colliding indices result in more than one insertion before an existing character (order irrelevant). The string length is now *M* ″ = *M* ′ + *n*_*ins*_
- *Padding or truncation*. If *M* ″ *> M*, truncate the string to length *M*. If *M* ″ *< M* pad the string to length *M* with random characters in {*a, c, g, t*}. The resulting string of length *M* is the garbled codeword. This padding/truncation implements the worst-case assumption that we have no independent information about where the true barcode begins or ends, but simply attempt to decode exactly *M* characters at the codeword’s nominal position (e.g., beginning of strand).

### 2.5 Fast Triage of Codewords by Trimer Similarity

We are committed to comparing *each* of *Q* (possibly many millions of) reads to *each* of *N* (possibly millions) codewords of length *M*, so as to find, for each read, that codeword with smallest Levenshtein distance. Naively, the number of implied operations is const ×*Q* × *N* × *M*^2^, where the constant is ∼10 and the factor *M*^2^ is the Levenshtein calculation. While feasible on a supercomputer, the implied∼10^16^ operation count is not to be recommended. Here, we show how to reduce it to∼10^13^ operations that can be done on a commodity GPU with 10^3^−10^4^ parallelism, implying as few as ∼10^9^ calculation steps, feasible on a single-head desktop machine.

We will employ a strategy of “triage”, that is, comparing each read to every codeword using only an approximate distance metric or similarity score, but one with a very small number of computer operations per comparison. This step will eliminate a large (often very large) fraction of possible identifications. Then, it is feasible to apply a more exact comparison to the small number of possible identifications that remain—either as a secondary triage (another approximate distance measure) or a true Levenshtein calculation. The final step will always be a Levenshtein (or similar) distance comparison, finding the decoding with the smallest true distance.

### Primary Triage by Trimer Hamming Popcount

Every codeword of length *M* has exactly *M* − 2 overlapping, consecutive trimers in {*a, c, g, t*}^3^ (a set of cardinality 4^3^ = 64). Why focus on trimers? Why not dimers or tetramers? The results of § 2.3 especially Figures 2 and S2 (right panels) suggest the need for barcodes of length *∼*30. On average the 4^2^ = 16 dimers will appear in a barcode about twice, and ≳ 85% will occur at least once. So there is relatively little information in either their uniqueness of occurrence or uniqueness of position. For tetramers, which number 4^4^ = 256, only ∼ 10% of them will appear in any given barcode, so *∼* 90% of the effort of keeping track of them is wasted. Trimers, each appearing on the order of once per barcode, is a unique sweet spot.

A suitable function *B*(*C*_*i*_) maps each codeword *C*_*i*_ to a 64-bit unsigned integer whose bits signify the presence (1) or absence (0) of each trimer in *C*_*i*_ The values *B*_*i*_ = *B*(*C*_*i*_) can be precomputed. Now, for each garbled *M* -mer read *R*_*j*_, we compute the distance measure

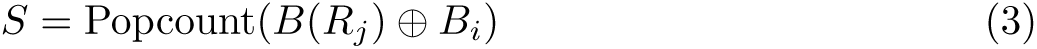

where ⊕ denotes bitwise exclusive-or and Popcount returns the number of set bits in a word, here the Hamming distance. Popcount is a single machine-language CUDA instruction on GPUs [39] that can readily be made accessible to PyTorch, or calculated (a few times slower) as a few-line PyTorch [40] function. Importantly, in either case, the calculation of equation 3 can be done in parallel across all *N* of the *B*_*i*_’s simultaneously. We may then eliminate from further consideration those codewords with the largest distances *S*, between 90% and 99%, depending on the DNA error rate (see further details below).

### Secondary Triage by Trimer Position Correlation

Conceptually, a secondary triage should need to be calculated, for each read, only for the list of codeword candidates that survive the primary triage. The output of the secondary triage would be an even shorter list of survivors. In practice, our proposed secondary triage is almost as fast as the above primary triage. That being the case, it is about equally efficient to apply the primary and secondary triages simultaneously to all the codewords, and then combine the triages, as will be described below. This strategy allows us to then jump directly to a Levenshtein comparison of the joint triage survivors.

Our secondary triage is motivated as follows: To be close in distance, a read *R*_*j*_ and codeword *C*_*i*_ should not only be similar by set-comparison of their trimers (popcount test above), but also close in the position indices, 0, …, *M −* 3, of identical trimers.

Denote individual trimers as *t ∈* [0, 64), and denote the ordered sequence of trimmers in a read or codeword as *t*_*i*_, *i* = 0, …, *M* − 2. Let *V* (*R*) be a function returning an integer vector of length 64, defined for a codeword or read *R* by the 64 components,

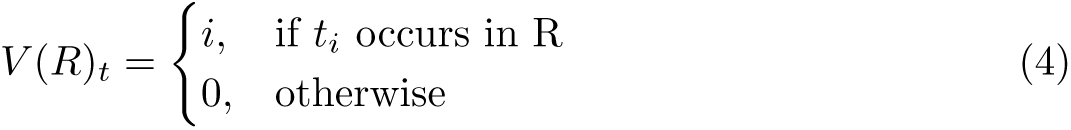

This is not quite a well-posed definition, because we might have *t*_*i*_ = *t*_*j*_ for *i* ≠ *j*, i.e., a collision in *V* (*R*)_*t*_. Supporting Information S4 discusses how collisions can be resolved in a computationally fast manner.

Now, the dot product *V* (*R*_*j*_) *· V* (*C*_*i*_), something like an unnormalized correlation of the two position functions, can be taken as a similarity measure. Since *V* (*R*_*j*_) and *V* (*C*_*i*_) can be precomputed, the dot products over all *i* and *j* can all be done in parallel on the GPU, exactly the kind of tensor calculation it is best at.

But why stop there? For any kernel function *K* that acts componentwise on a vector, the dot product *K*(*V* (*R*_*j*_)) · *K*(*V* (*C*_*i*_)) is also a similarity measure. Multiple *K*’s return different similarity information. We find that kernels of cosine shape,

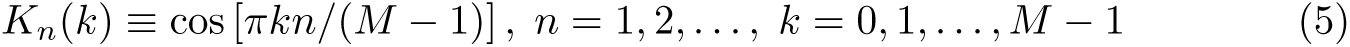

(with small *n*), each give good results, even better when combined as next described. The intuition here is that values *n >* 1 are more sensitive to the ordering of trimers on finer scales, but only up to some value of *n* where indels result, on average, in a loss of phase coherence with the cosines. We find that 1 ≤ *n* ≤ 4 works well, with larger values giving little improvement.

### Combined Triage

Although operations across all pairs of reads and codewords are by definition expensive, we have found it efficient to expend the cost of ranking (i.e., sorting) each read’s *N* distance scores (against every codeword) for the handful of distance measures, equations 3 and 5 (with *n* = 1, 2, 3, 4). Let *r*(*i, n*) denote the rank of the *i*th codeword in the *n*th distance measure, small ranks meaning most similar. Then we define the combined distance measure *r*(*i*) as the product

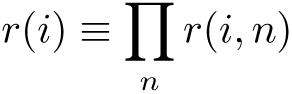

This can be viewed as akin to a naive Bayes estimate, since *r*(*i, n*)*/N* is something like a Bayes evidence factor provided by the distance measure *n*. Finally, we rank the *r*(*i*)s.

Figure 3 shows results for a simulation with *N* = 10^6^ codewords of length *M* = 34 whose reads are corrupted (using the error model describe above) with

**Figure 3.**
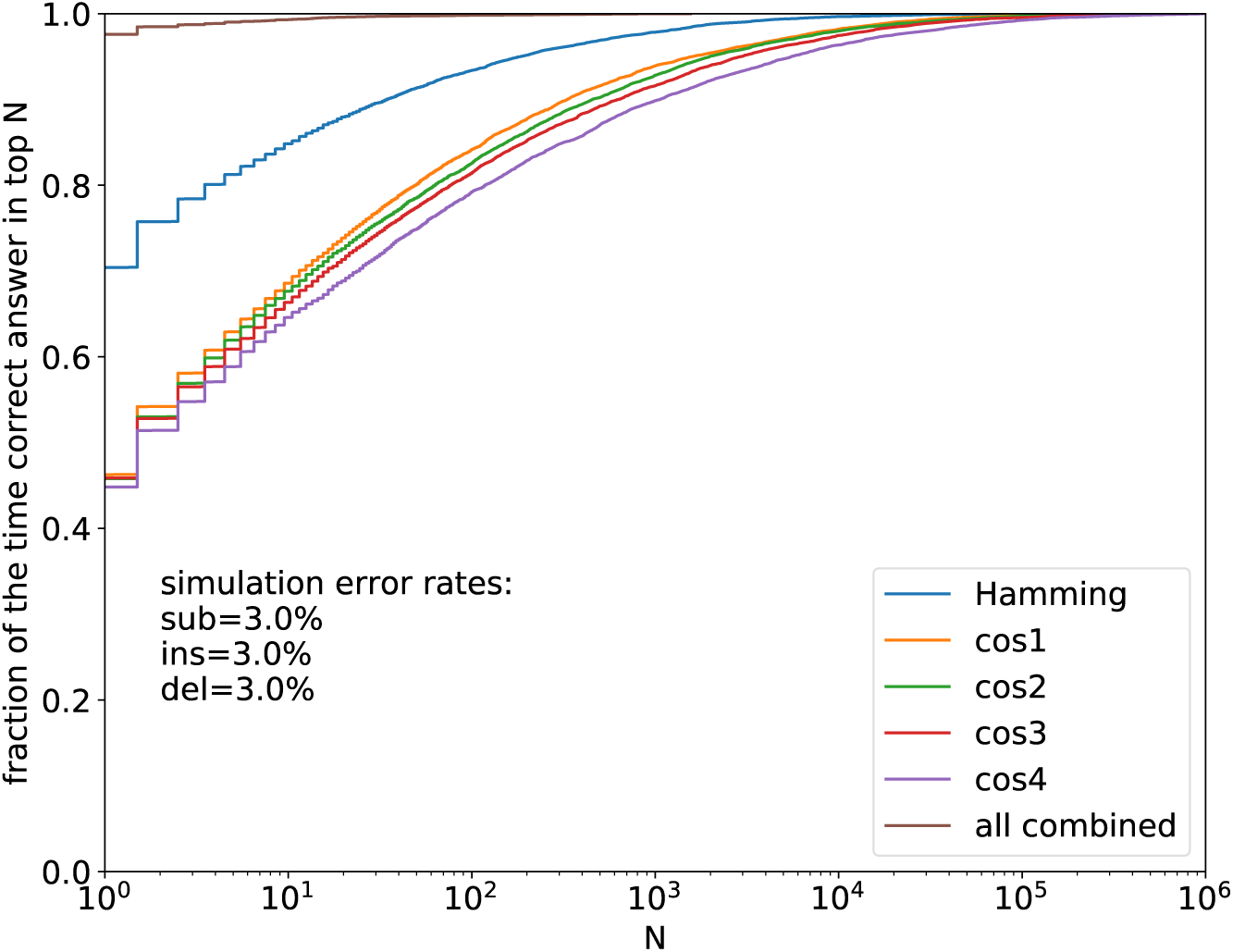
Triage performance of individual filters and combined. For *N* = 10^6^ codewords of length *M* = 34 nt, and for an error model with *p*_*sub*_ = *p*_*ins*_ = *p*_*del*_ = 0.03, the figure shows the probability of capturing the correct codeword with triage to sets of codeword possibilities much smaller than *N*. Here, after triage and with negligible loss of recall, exact Levenshtein testing is needed for fewer than 10^3^ codewords.

*p*_*sub*_ = *p*_*ins*_ = *p*_*del*_ = 0.03, a total error rate of 9%. One sees that, here, the Hamming popcount is doing most of the heavy lifting, but combining with position similarity gives substantial improvement. In the figure “cos*n*” denotes the kernel functions in equation 5 While these have very similar performance individually, the elimination of any of them decreases the combined performance somewhat.

In this example, triage from 10^6^ down to 10^3^ codeword possibilities for each read would capture the correct answer almost always (*≫* 99%) so that the exact Levenshtein calculation could be done on only the smaller set, at negligible computational cost.

A somewhat less favorable, but still very feasible, example is for the large error rates *p*_*sub*_ = 0.05, *p*_*ins*_ = 0.05, *p*_*del*_ = 0.10, as shown in Figure 4. Here, triage from 10^6^ to 10^5^ produces negligible loss of recall. We will see in Results that parallel computation of Levenshtein on 10^5^ codewords per read is also very feasible.

**Figure 4.**
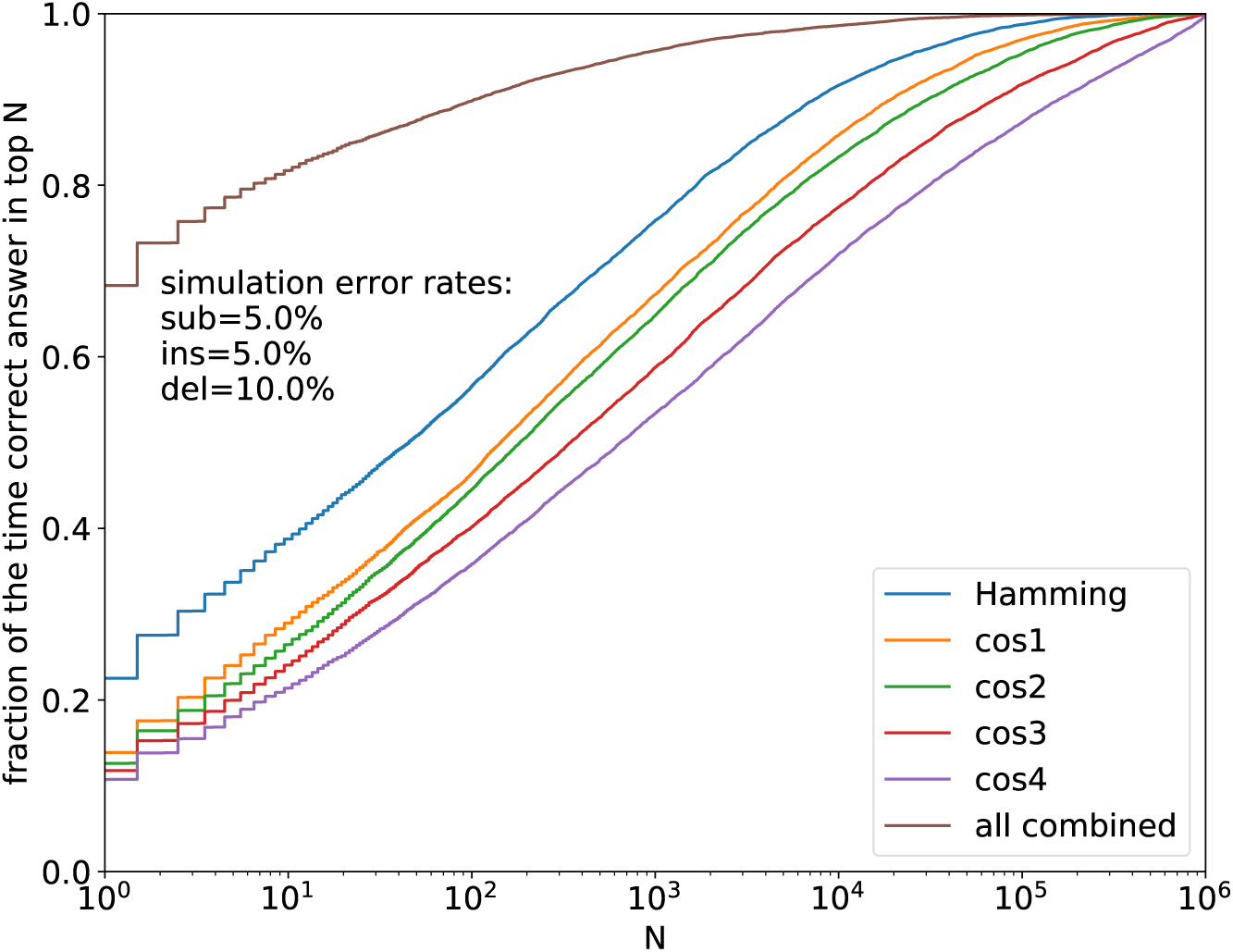
Same as Figure 3, but with error rates *p*_*sub*_ = 0.05, *p*_*ins*_ = 0.05, *p*_*del*_ = 0.10. Here, further testing on about a tenth of codewords is required.

**Figure 5.**
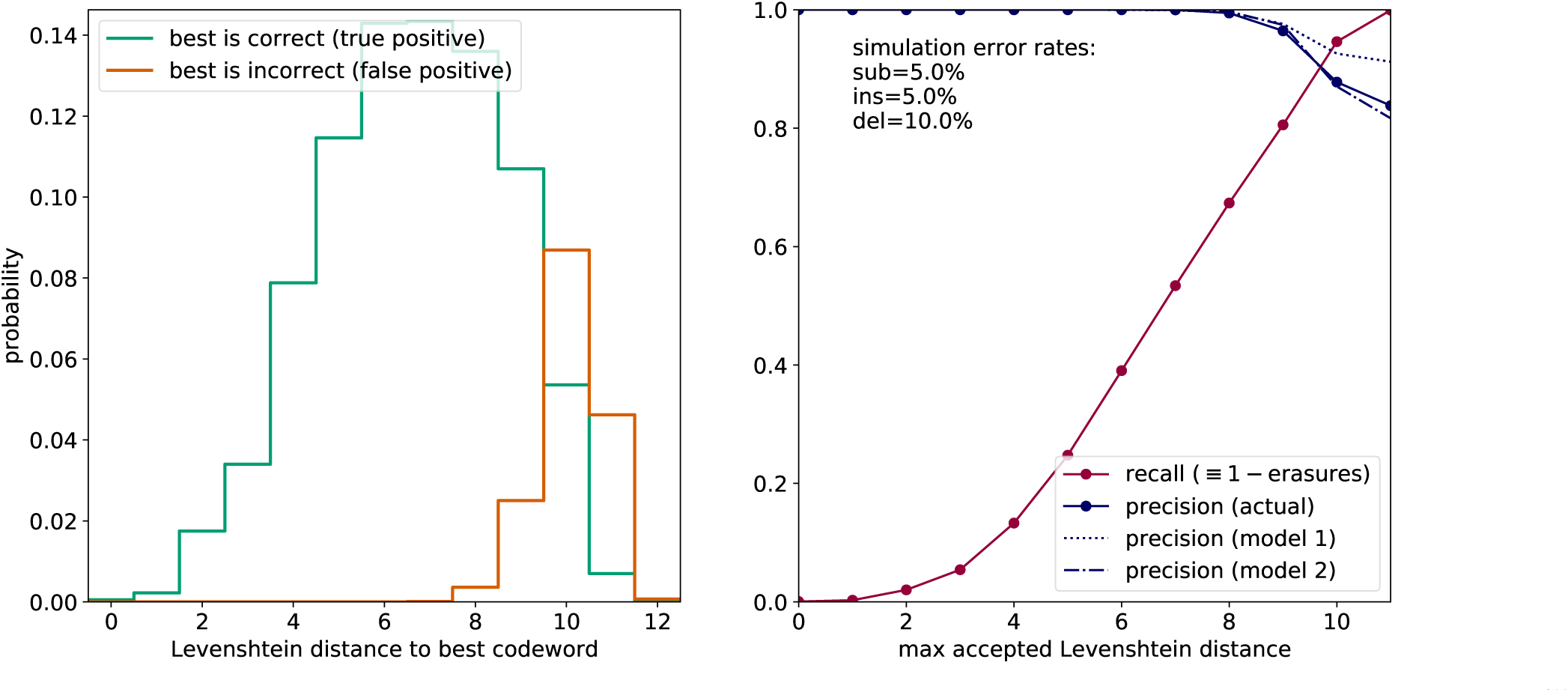
Simulation with 20% DNA error rate. Left: Distribution of distances seen when decoding to the Levenshtein-closest codeword among 10^6^ possibilities. The experimenter, not knowing which decodes are correct, sees the sum of the red and green histograms. Right: With data from the left panel, for each choice of threshold *T*, recall is the fraction of all events (green plus red) ≤ *T*. This is knowable to the experimenter. Precision is, for events ≤ *T*, the fraction of green (versus red) events. This is not directly knowable but can be estimated by the models shown (see text).

## 3 Results

Illustrating the use and practicality of the above methods, we here give the results of detailed simulations for the case of one million barcodes (*N* = 10^6^) of length *M* = 34 nucleotides in the presence of end-to-end DNA total error rates of 20% per nucleotide (base case) with excursions to smaller (9%) and larger (30%) rates. These rates are intentionally chosen to be all very large as compared to next-generation NGS error rates, and even large or very large as compared to third-generation TGS rates (see Section 1). We know of no previously proposed barcode sets capable of success with these parameters at plausible computational workloads.

### 3.1 Precision and Recall

It is important to emphasize that the methods of this paper do not give either perfect precision, that is, the correct decoding of *every* garbled read independent of its number of errors; nor perfect recall, that is, no garbled reads rejected as undecodable erasures. Rather, by choice of an integer threshold Levenshtein distance *T*, the user may set any desired recall between 0% (all erasures) and 100% (no erasures) and must then accept the implied level of precision.

For these tests, we generated either random sets of codewords, or else otherwise random sets that excluded codewords with homopolymer runs of *>* 3, or CG or AT fraction greater than 0.66. There was no discernible difference in results between same-sized fully random and sequence-constrained codeword sets.

Garbled reads were assigned to the closest codeword by Levenshtein distance when the distance was≤ *T*, otherwise called as erasures. For the base case of 20% total errors, with *p*_*sub*_ = 0.05, *p*_*ins*_ = 0.05, *p*_*del*_ = 0.10, the figure shows results for the full range of choices of *T*. In a real experiment, the user does not know which decodes are correct, so sees the sum of true and false positives (green and red histograms). That is enough to calculate the recall for each possible value of *T*, but not the precision, which requires “knowing the answers”.

However, the user does know that there is *some* red histogram whose expected shape was already calculated above in equations 1 and 2 and Figure 2. The user also knows that the green histogram should be roughly binomial, but “censored” by the red histogram in a computable way. In Supporting Information S5, we show that this is enough information to model the expected precision function either naively (shown as Model 1) or, with additional assumptions, somewhat more accurately (shown as Model 2). So, in practice, the user can use these models to choose an appropriate value *T*. In the figure, a suitable choice based only on the models might be *T* = 8, which in simulation gives 99.6% precision with 67% recall. Whether this is sufficiently large recall to be useful depends on the design of the experiment, for example, whether a given barcode is expected to be read several or many times, in which case a 33% loss to erasures can be tolerable.

Supporting Information S6 shows the analogous figures for error rates 10% (with *p*_*sub*_ = 0.033, *p*_*ins*_ = 0.033, *p*_*del*_ = 0.033) and 30% (with *p*_*sub*_ = 0.075, *p*_*ins*_ = 0.075, *p*_*del*_ = 0.15). For the former of these, the choice *T* = 9 yields precision 99.9% with recall 98.8%. For the latter, recall must be sacrificed to get good precision. *T* = 7 gives precision 99.8% with recall 20.4%, while *T* = 8 gives precision 98.2% with recall 32.6%. The user is assumed to know something about the experimental DNA error rate a priori. However, if this is not the case, then the the above values can be assumed as lower bounds. Specifically, for any assumed total error rate significantly less than 30%, the value *T* = 8 should give *>* 99% precision along with a recall that will be immediately known from the data, by the number of erasures called.

### 3.2 Performance and Cost

The minimum requirement for using the methods described in this paper is a compute node (or cloud instance) with at least 2 CPU cores and at least 1 commodity- or server-grade GPU having at least 8 GB memory. To use exactly our code (as available on Github), PyTorch [40] and its associated software tool chain is required, but porting to other CUDA tool chains (e.g., TensorFlow [41] should be straightforward.

We measured actual performance on a standalone workstation with an Intel i9-10900 processor (10 CPU cores, 20 logical processors) and 2 Nvidia RTX 3090 GPUs, each with 10,496 CUDA cores and 328 tensor cores. The purchase price of this machine (year 2022) was US$12,000. The (year 2022) marginal cost of adding additional comparable GPUs would be about US$1,500 each.

As a typical performance test, we generated 1,000,000 simulated reads of 34 nt random barcodes with 20% error rates. (Performance does not actually depend on error rate.) Reads were divided among processes running concurrently on separate CPU cores. On the above machine, we found that four such processes, two assigned specifically to each GPU, gave best performance, saturating the two GPUs (and four CPUs) at close to 100% usage. Memory usage per GPU was 7.4 GB. Wallclock execution time was 4943 seconds, implying about 17.5 million reads per 24-hour day. This is likely adequate performance for many applications and will only improve with time as GPU cycles get faster and cheaper.

For applications requiring greater throughput, there are various options: Academic supercomputer centers allocate time (at zero cost) competitively to academic users. A current example is the Longhorn computer at the Texas Advanced Computing Center (TACC) [42] with 384 Nvidia V100 GPUs, implying on the order of 7 billion reads per day for the full machine. The “startup” allocation of 100 node hours should process on the order of 150 million reads, and much larger allocations are routinely awarded. Alternatively, commercial cloud instances of GPUs can be stood up by the hour in any desired quantities and thus any desired throughput. Current (2022 [31],[32]) prices of about US$ 0.50 per GPU-hour imply a cost of about US$ 1.50 per million reads processed. This can be compared to the 3-year amortized cost of the standalone machine described above implying about US$ 0.60 per million reads; or the amortized marginal cost of each additional GPU, which implies about US$ 0.15 per million reads.

## 4 Discussion

The main point of this paper is demonstrating the practicality of all-to-all comparisons for closest Levenshtein or Needleman-Wunsch match (that is, comparing all reads to all barcodes) with DNA barcodes sets of 10^6^ barcodes or larger, and for reads numbering many millions or more. The elements that make this possible are (1) the parallel processing capabilities of current commodity GPUs, (2) the use of a novel, very fast, parallel triage that, for each read, eliminates from competition all but a small fraction of candidate barcodes, and (3) the ability to parallelize the Levenshtein or Needleman-Wunsch computation a significant degree, both within a single calculation and across many such.

All-to-all comparison in turn makes practical the use of random barcode sets (defined and fixed for each experiment) that derive error-correcting capability simply by the statistics of their average distances from one another. While the required lengths, ∼ 30 nucleotides, may be undesirably long for use with short read lengths, they are not a detriment with read lengths of third-generation sequencing. And, in that context, the ability to use of direct, parallel Levenshtein (or an even faster approximation as discussed), allows as many as 6 to 8 errors to be corrected (set above by the threshold value *T*), along with correctly flagging as undecodable “erasures” reads with more than this number. At 10% errors per nucleotide, considered a large value, we are able to demonstrate precision of 99.9% and recall of 98.8%. Even with 20% errors per nucleotide, we demonstrate 99.6% precision with recall of 67% (meaning that at most 1/3 of reads are wasted). We know of no other existing, practical DNA barcode methodologies that are able to operate in these high-error-rate regimes with ≳ 10^6^ barcodes. In these statistics, errors in barcode synthesis (“wrong barcodes”), are as equally correctable as errors created at later stages of an experiment or during final sequencing.

In contrast to this paper, mathematically constructed error correcting codes (ECCs) of a given length *L* are designed to have fewer near-collisions than our random barcodes of the same length. If there existed known mathematical ECCs capable of (i) correcting as many as 6 to 8 errors, and (ii) correcting not just substitution errors, but also insertions and deletions (indels), then these would be superior to random barcodes. But we know of no such ECCs. [27]

## Code Availability

Python and PyTorch code for all the computations in this paper are available on Github at https://github.com/whpress/RandomBarcodes.

## Acknowledgments

I thank John Hawkins for valuable discussions, and thank Anshuman Suri for contributing a useful piece of CUDA coding (providing PyTorch access to the CUDA popcount primitive). Ilya Finkelstein and Jim Rybarski made useful comments on the manuscript.

## Supporting Information

### S1. Parallel Computation of Needleman-Wunsch and Levenshtein Distances

These distances are conventionally calculated by dynamic programming on a Cartesian tableau, filling in squares from top to bottom and left to right (see Wikipedia, “Needleman-Wunsch algorithm”). Mapping this to the CUDA strided-slice tensor programming model on a GPU is facilitated by tilting the tableau 45°, thus displaying the desired calculation as a parallel calculation on a top-to-bottom directed acyclic graph, shown as the red grid in Figure S1. Pink dots on the blue grid are filled with multiples of the skew penalty. The Boolean tensor [*A*_*i*_ == *B*_*j*_] is calculated in parallel on the red grid, then mapped to the blue. The calculation then proceeds top to bottom on the blue grid filling in the green dots. Some parallelism is generated by doing each row’s green dots in parallel (shown in the Figure as at most three, but actually as many as *M* − 1 for *M* nt codewords. A much larger parallelism is achieved by “stacking” 10,000 to 50,000 such grids out of the plane, each with the same string A (shown as *A*1, *A*2, *A*3) but different strings B (shown as *B*1, *B*2, *B*3, *B*4). PyTorch code for this algorithm is included on GitHub.

**Figure S1.**
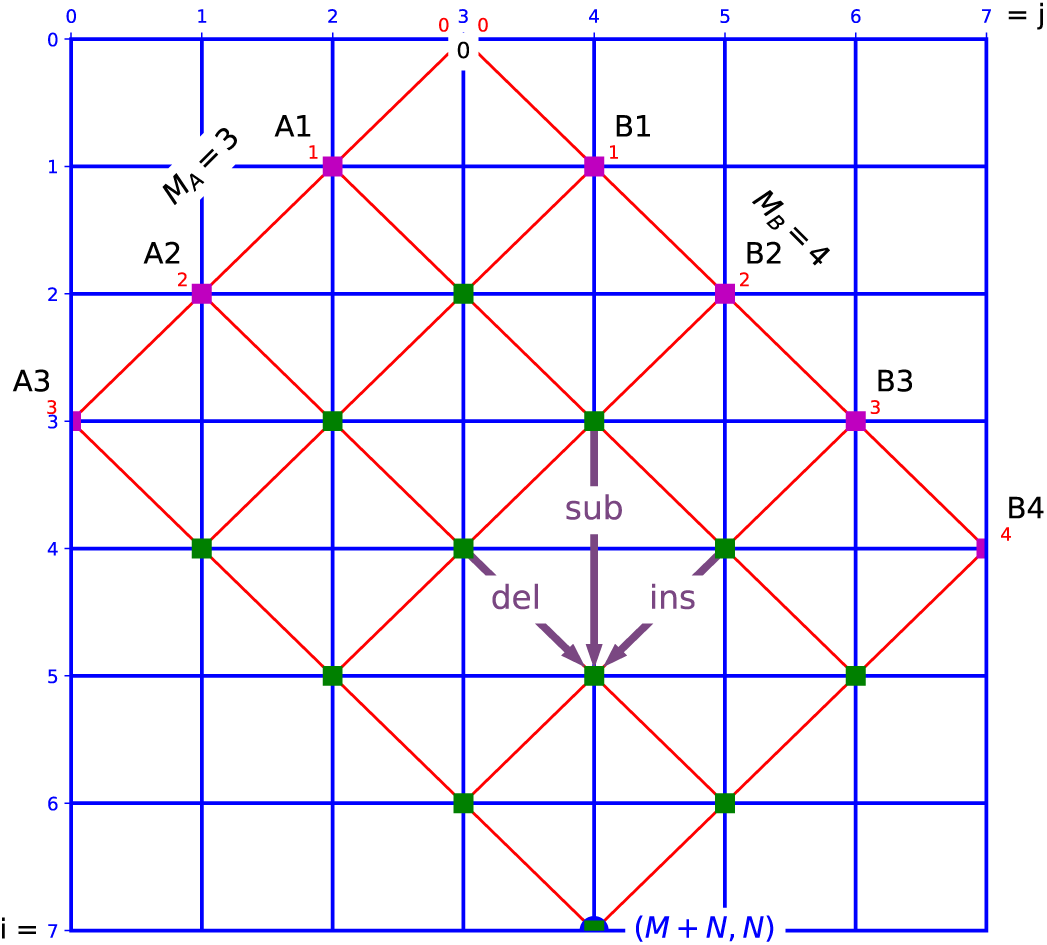
One-to-many parallel Needleman-Wunsch or Levenshtein distance calculations. The red grid is largely conceptual. The main calculation is done in parallel on a stack of many blue grids, from top to bottom in the figure. Green dot values are filled with the minimum among del, sub, and ins, i.e., the shortest path to that dot. The stack of answers appears as the blue dot at the bottom.

#### Approximate Levenshtein

Referring to the red tableau in the figure, and to the sub, ins, del arrows, a parallel calculation can be done for substitutions and insertions a full (red) row at a time, but not for deletions, because the value in an earlier column can affect a later one. Alternatively, a fully parallel calculation a column at a time is possible for substitutions and deletions, but not insertions.

That said, we can ask what happens if we ignore this reality and just do the fully parallel calculation? The answer will be wrong only slightly and only when there are two or more consecutive deletions (if processing by rows) or insertions (if processing by columns). Moreover, if we do the parallel deletion (insertion) step twice, literally repeating the same one line of code, then the answer will be wrong only when there are *three* or more consecutive deletions (insertions), a relatively rare occurrence.

This parallel “approximate Levenshtein” calculation is found to be several times faster on a commodity GPU than the parallel exact Levenshtein calculation. Approximate Levenshtein can be used as a “tertiary triage” (in the language of section 2.5), bringing the computational burden of exact Levenshtein down to almost negligible. In fact, in the simulations that we have tried, the use of approximate Levenshtein alone, without any other followup, gives results for recall and precision (section 3.1) that are virtually indistinguishable from those where exact Levenshtein is used. Since the goal is correct decodes, not exact Levenshtein distances, the use of approximate Levenshtein seems justified by the performance gain.

### S2. Fitted Polynomial Expression for Levenshtein Distance Distribution of Random Strings

Figure 1 in the main text indicated by dots the values actually obtained by simulation, which have probabilities as small as ≲ 10^*−*7^. For smaller probabilities, we need to extrapolate. Rather than fit each value *M* separately, which would allow extrapolation on Levenshtein distance *L*, but not interpolation on codelengths *M*, we fit a bivariate polynomial,

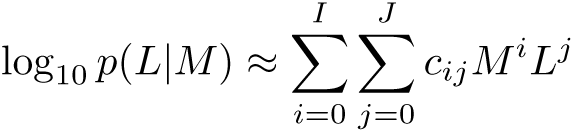

Here *p*(*L*|*M*) is the probability that two random *M* nucleotide strings are separated by a Levenshtein distance *L*. Bivariate fitting also acts to improve the accuracy, because an improbably deviant small sample in the tail of one *M* value is mitigated by the other *M* values.

The coefficients *c*_*ij*_ for the adopted best fit are,

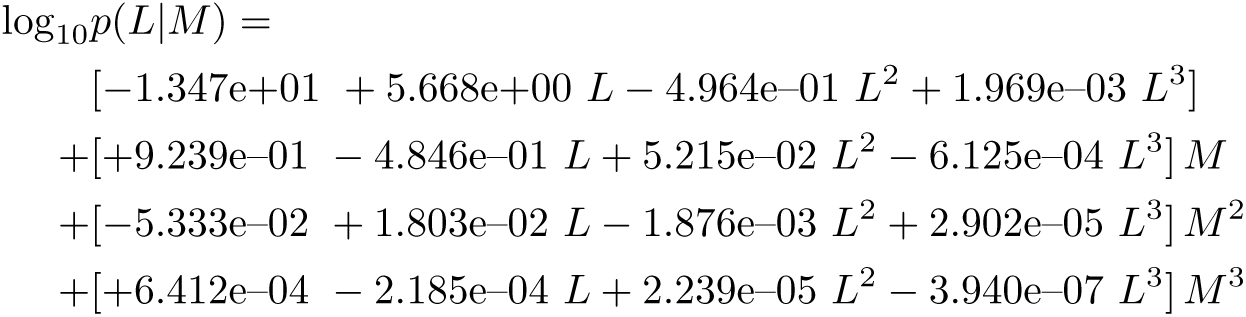

### S3. Distribution of Closest Non-Causal Distance to a Set of *N* Codewords for Other Nucleotide Lengths

See main text Figure 2, which was for the case of *M* = 34-mer codewords. Here are analogous figures for *M* = 30, 26, 22, and 18.

**Figure S2.**
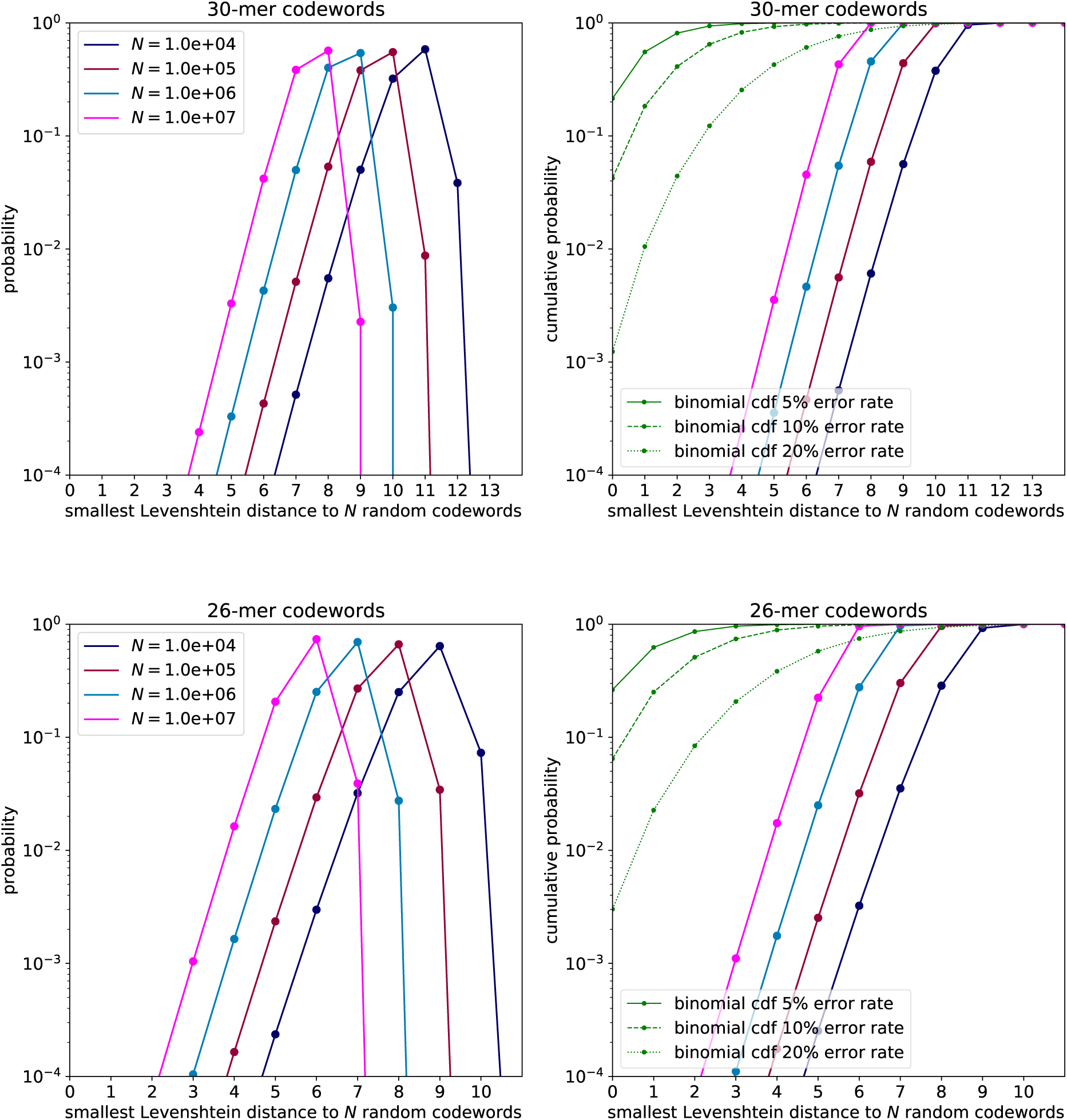
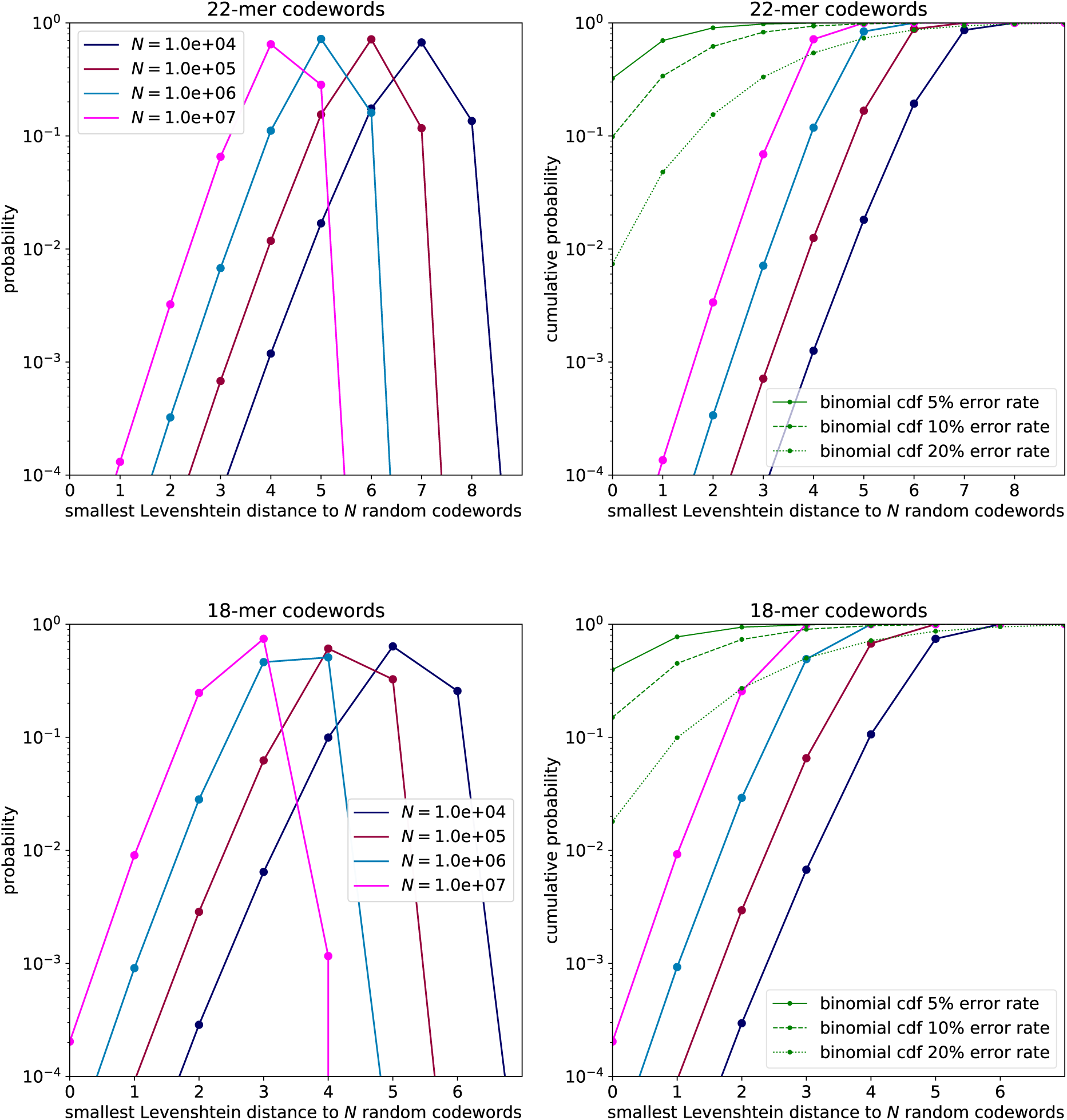
More Smallest Levenshtein Distances to Random Sets of Codewords.

### S4. Collisions in the Trimer Position Function

We noted above that equation 4 might not define a single-valued equation for the positions of each trimer, because a specific trimer (“cgt” for example) might occur in more than one position. In practice, it is not too bad to pick any one position, randomly. Such a function *V* (*R*)_*i*_ returns a (any) position in the codeword for each of 64 trimers *t*_*i*_, defining the position to be zero if the trimer does not occur in the read *R*.

Even better results are obtained by combining all positions {*i}* of a given trimer *t* causally into some kind of pseudo-position. This can be done either before or after applying the kernel *K*() in equation 5. After some trial and error among alternatives, we replace the components of the vector *K*(*V* (*R*_*j*_)), that appears in the dot product, by

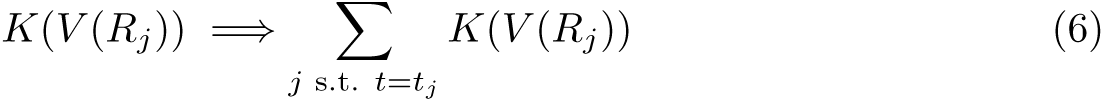

that is to say, we sum the colliding kernels (in our case, cosines) before taking the dot product. The sums can be done in parallel by a scatter-add operation.

### S5. Models for Estimating Precision

Here we develop two models that allow a user to estimate, for codewords of length *M*, as a function of threshold Levenshtein distance *T*, the precision of decoded garbled barcode reads. The models can then be used to choose a value *T* that appropriately trades off precision and recall. We assume that the user has an estimate for the total error rate *r* (or chooses some value as an upper bound).

#### Model 1

For every read, there is a distribution of Levenshtein distances *L*_*t*_ from its true (causal) codeword that we model as a binomial probability binom(*L*_*t*_|*M, r*); and there is a distribution *P* (*L*_*f*_) of its distances *L*_*f*_ from the closest false (non-causal) codeword, which was calculated above in Section 2.3. When we have *L*_*t*_ ≤ *T* and *L*_*t*_ *< L*_*f*_ we can score a true positive (TP). For ties *L*_*t*_ ≤ *T* and *L*_*t*_ = *L*_*f*_, we resolve the tie randomly and score half a true positive. Conversely, when we have *L*_*f*_ ≤ *T* and *L*_*f*_ *< L*_*t*_ we can score a false positive (FP), or half a false positive if *L*_*f*_ = *L*_*t*_. The remaining case is when *L*_*t*_ *> T* and *L*_*f*_ > *T*, which is an erasure.

Parsing these inequalities in the two-dimensional grid of *L*_*t*_ and *L*_*f*_, and with the assumed probability distributions, gives the rates for TP and FP,

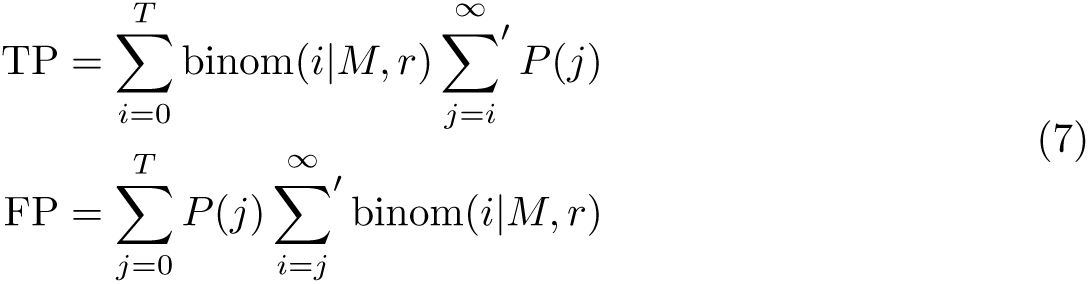

Here Σ′ denotes a sum with a factor 1/2 applied to its first term. In terms of these quantities,

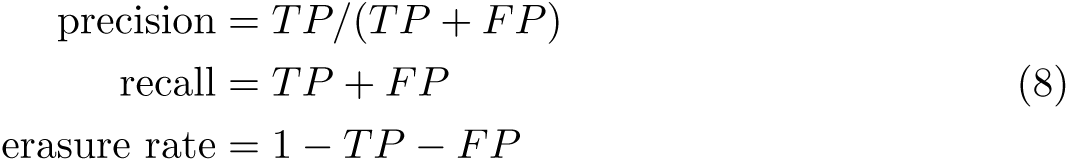

#### Model 2

A weakness is Model 1 is its assumption of a binomial distribution for *L*_*t*_, when we know this is not correct with indels. Also, Model 1 does not make use of the experimentally observable distribution of distances *L*, a mixture of the causal and non-causal distributions,

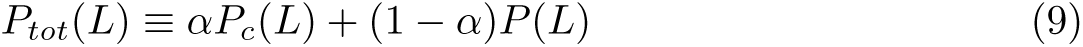

(In Figure 4, this mixture was shown as the green and red histograms.) Implicitly, Model 1 estimated *P*_*c*_(*L*) by

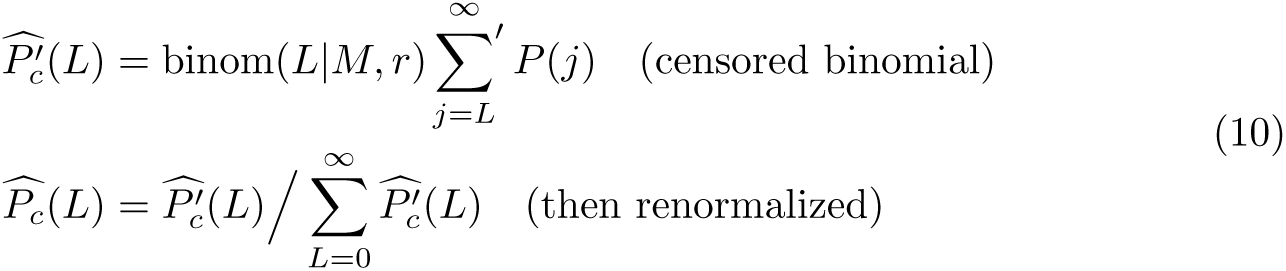

where “hat” denotes an estimator.

For Model 2, we first least-squares estimate *α* in equation 9 by the formula,

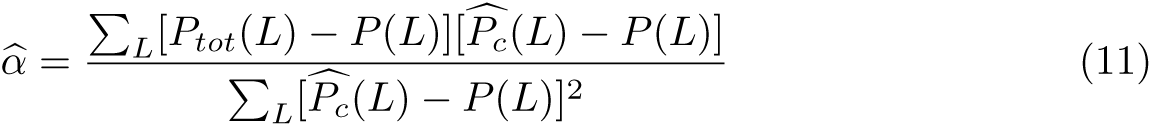

in terms of which the precision is then estimated by

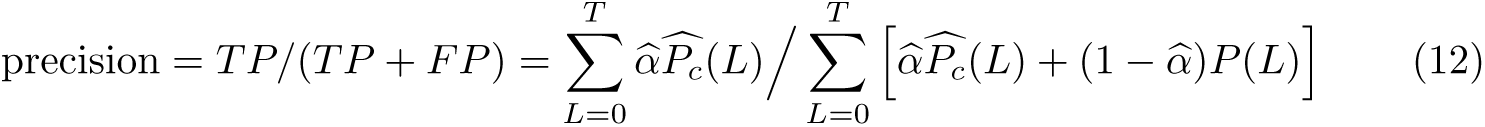

It is not obvious mathematically that Model 2’s precision estimate must be better than Model 1’s, but in simulation we generally find it to be.

### S6. Precision and Recall for Other Simulated DNA Error Rates

These figures are analogous to Figure 3, but for the different DNA error rates 10% and 30%.

**Figure S3.**
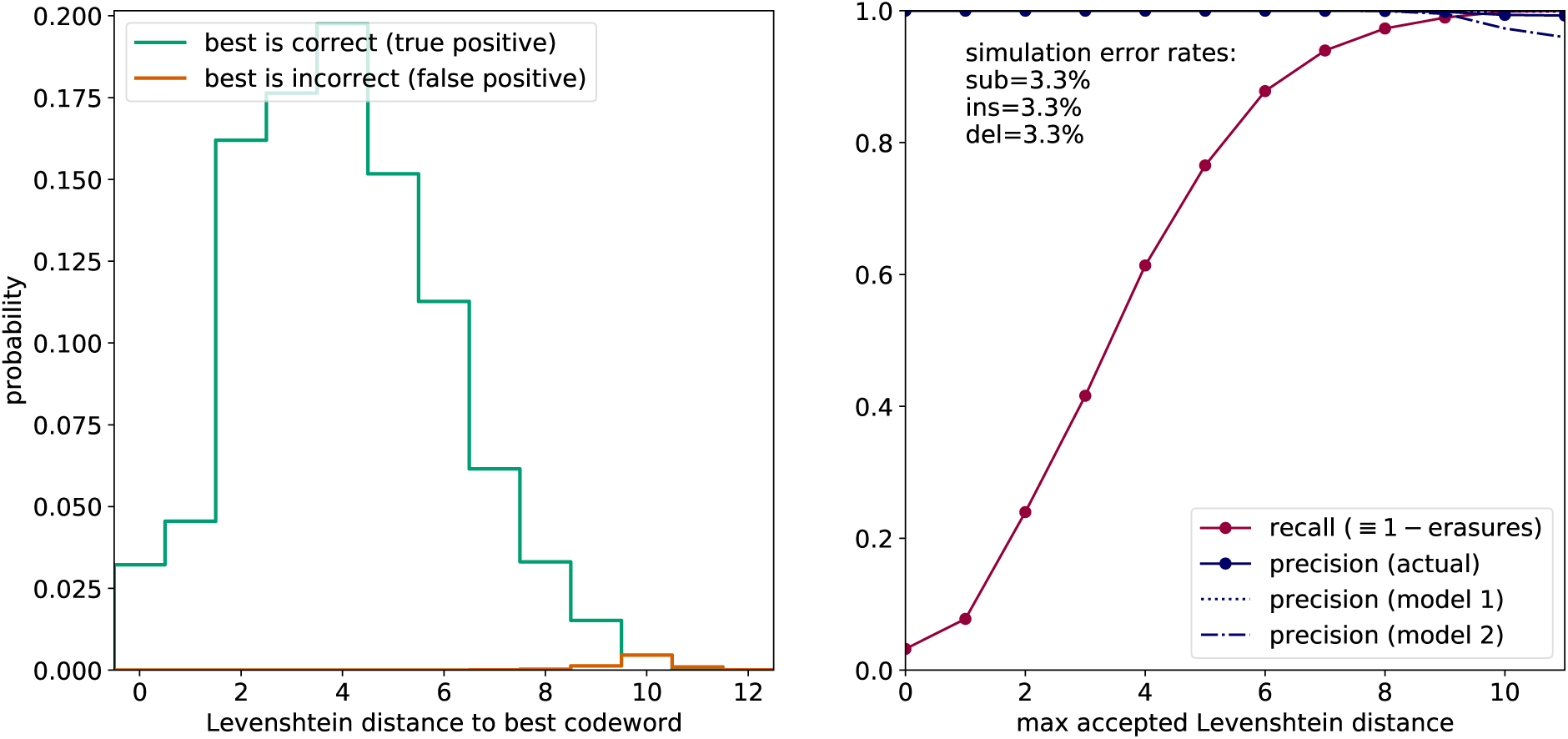
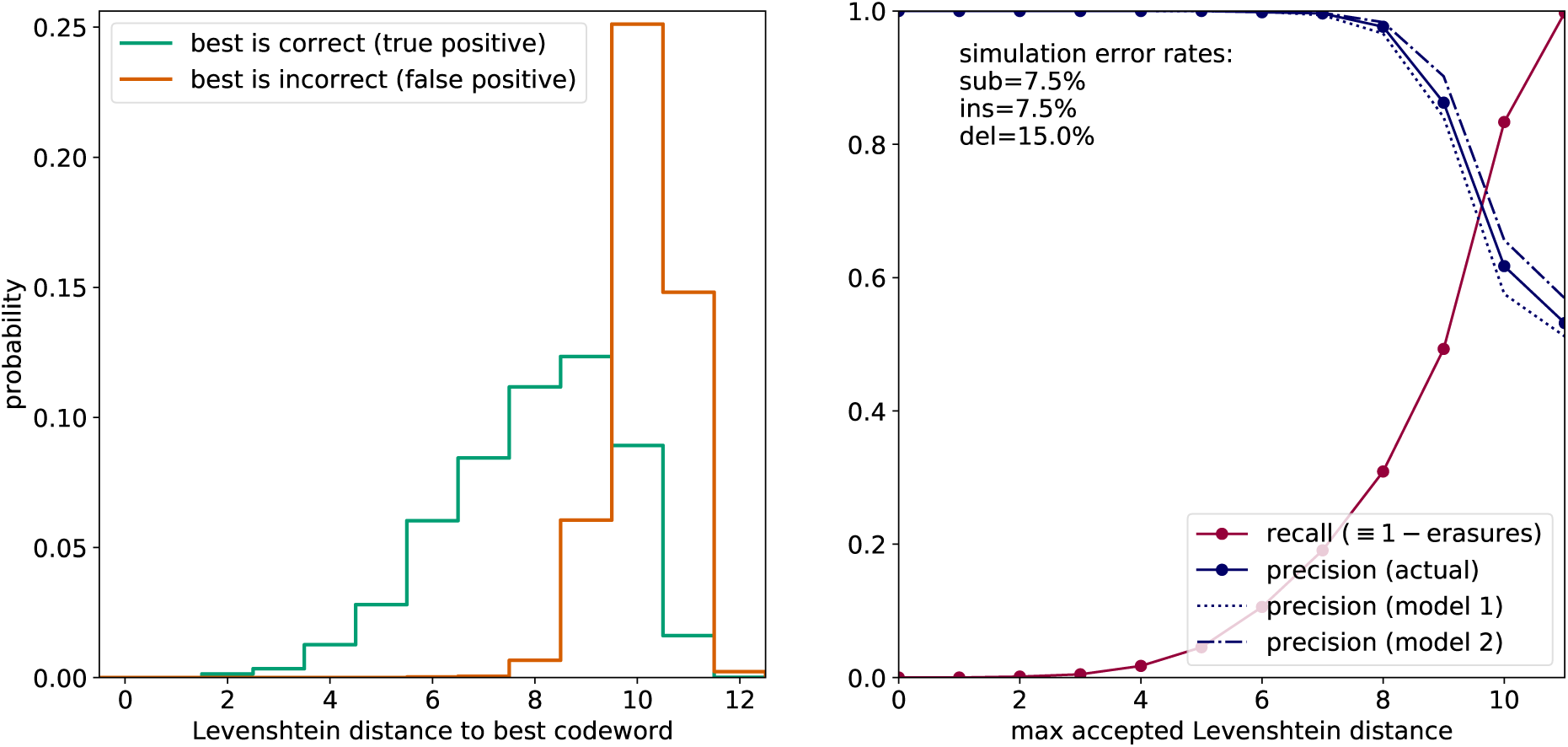
Precision and Recall Simulations for Additional DNA Error Rates.

